# Pathologist-interpretable breast cancer subtyping and stratification from AI-inferred nuclear features

**DOI:** 10.1101/2025.09.04.674077

**Authors:** Ranjan Kumar Barman, Saugato Rahman Dhruba, Danh-Tai Hoang, Eldad D. Shulman, Emma M. Campagnolo, Andy T. Wang, Stephanie A. Harmon, Tom C. Hu, Antonios Papanicolau-Sengos, MacLean P. Nasrallah, Kenneth D. Aldape, Eytan Ruppin

## Abstract

Artificial intelligence (AI) is making notable advances in digital pathology but faces challenges in human interpretability. Here we introduce **EXPAND** (<u>EX</u>plainable <u>P</u>athologist <u>A</u>ligned <u>N</u>uclear <u>D</u>iscriminator), the first *pathologist-interpretable* AI model to predict breast cancer tumor subtypes and patient survival. EXPAND focuses on a core set of 12 *nuclear pathologist-interpretable features* (*NPIFs*), composing the Nottingham grading criteria used by the pathologists. It is a fully automated, end-to-end diagnostic workflow, which automatically extracts NPIFs given a patient tumor slide and uses them to predict tumor subtype and survival. EXPAND’s performance is comparable to that of existing deep learning non-interpretable black box AI models. It achieves areas under the ROC curve (AUC) values of 0.73, 0.79 and 0.75 for predicting HER2+, HR+ and TNBC tumor subtypes, respectively, matching the performance of proprietary models that rely on substantially larger and more complex interpretable feature sets. The 12 NPIFs demonstrate strong and independent prognostic value for patient survival, underscoring their potential as biologically grounded, interpretable biomarkers for survival stratification in BC. These results lay the basis for building interpretable AI diagnostic models in other cancer indications. The complete end-to-end pipeline is made publicly available *via* GitHub (https://github.com/ruppinlab/EXPAND) to support community use and reproducibility.

## INTRODUCTION

Breast cancer (BC) is the most common women’s malignancy around the world with significant mortality and morbidity rates^1,2^. Pathological diagnosis is pivotal in guiding clinical management and optimizing treatment strategies for such patients^2^. This process begins with the pathologist carefully examining patient histopathology images, primarily obtained from the hematoxylin and eosin (H&E)-stained tissue slides, to identify relevant patterns leading to the diagnosis^3,4^. Recent breakthroughs in AI, particularly in deep learning (DL), and digital pathology are revolutionizing cancer diagnostics and prognosis by unlocking the ability to rapidly and precisely detecting key patterns in H&E-stained whole-slide images (WSIs), enabling more precise and personalized patient care^5–10^. AI in digital pathology offer tremendous potential for advancing precision medicine by uncovering novel biomarkers^11–14^, predicting treatment response^15–18^ and integrating multi-omics data^19–22^, beyond its immediate clinical applications^23–25^.

While deep learning has significantly advanced precision oncology, its approaches are frequently seen as lacking clarity and interpretability, resulting in the common characterization of these models as “black boxes”. These models can automatically learn highly informative but rarely intuitive features from raw images which yield high accuracy but obscure the decision-making process, raising concerns about their applicability in routine clinical practice^26–29^. Addressing the interpretability challenge is crucial for building trust and ensuring the effective integration of AI-driven insights into precision oncology^30,31^. Two prominent approaches have emerged to address the challenge of developing such interpretable and explainable AI models in cancer diagnostics and prognosis: (1) A *top-down approach* that leverages the latest developments in the attention-based transformer architecture to offer an intuitive but indirect form of interpretability *via* highlighting the informative regions for prediction in the H&E-stained WSIs, but leaves much of the underlying decision-making process hidden^9,32–37^; and (2) A *bottom-up approach* that extracts explainable features from these H&E-stained slides, such as the human-interpretable image features (HIFs)^38^, that can be directly applied for prediction, yielding a more transparent solution. The bottom-up approach, while often offering lower predictive power than the top-down approach (reflecting the typical trade-off between accuracy and interpretability in AI), aligns closely with the established pathology workflow. Thus, potentially making it more user friendly for clinical applications in pathologist workflow^38–40^.

Few recent efforts have focused on developing HIFs-based methods to predict clinical outcomes in cancer *e.g.*, tumor subtype, genomic instability, patient survival and so on^38–40^. Amgad *et al.* obtained 109 HIFs from H&E slide images to develop the Histomic Prognostic Signature (HiPS) that can successfully predict survival risk in BC, demonstrating the potential of HIFs to aid in clinical decision-making^40^. Diao *et al.* extracted 607 HIFs through a DL-based approach and further demonstrated their ability to predict clinically relevant molecular phenotypes in cancer, highlighting their role as a bridge between traditional pathology and modern AI techniques^38^. Abel *et al.* later identified 90 nuclear morphological features *i.e.*, nuclear HIFs (nuHIFs), and demonstrated their effectiveness in detecting genomic instability across three cancer types and BC molecular subtypes, underscoring their potential in precision oncology^39^.

While the HIFs-based methods show great promise, the large number of features—though labeled as “human-interpretable” do not fully reflect the pathology-based decision-making process. Despite their utility, many of the features do not align with the way pathologists assess tissue morphology, highlighting the need for a more streamlined and intuitive approach to truly address the interpretability challenge. To this end, we aim to develop the first truly *pathologist-interpretable AI model* that predicts patient tumor subtypes and survival as accurately as currently possible by deep learning non-interpretable models while providing clear, transparent insights on how it arrives at these predictions.

Building on the recent studies reviewed above, we first leveraged the 607 HIFs and 90 nuHIFs, derived from cell counts and tissue region properties, to develop a set of 25 **pathologist-interpretable features (PIFs)** and later converged to 12 **nuclear PIFs (NPIFs)**, offering greater interpretability. To ensure broad applicability, we design an end-to-end pipeline called **EXPAND** (<u>EX</u>plainable <u>P</u>athologist <u>A</u>ligned <u>N</u>uclear <u>D</u>iscriminator) that obtains these 12 NPIFs from patient WSIs (either selects from pre-extracted HIFs or extracts *via* nuclear segmentation) and builds interpretable models to predict their tumor subtypes and corresponding survival risks. Our approach provides high performance as currently possible in clinically relevant applications while maintaining high levels of interpretability, making it ideal for aiding pathologists and clinicians for both routine and critical diagnostic and prognostic tasks, reinforcing the value AI in precision oncology.

## RESULTS

### Overview of the study

EXPAND is a bottom-up modeling approach based on the PIFs extracted from patient H&E-stained whole-slide images to ensure interpretability and applicability to clinical tasks. We first compiled two recent sets of interpretable features: 607 HIFs and 90 nuHIFs outlined by Diao *et al.* and Abel *et al.*, respectively^38,39^. Diao *et al.* trained two convolutional neural networks (CNNs) using pathologist-annotated WSIs to detect cell and tissue types, enabling the computation of 607 HIFs that describe cell densities, tissue composition and how cells are arranged relative to each other^38^. Abel *et al.* followed it up by extending the framework to nuclear morphology through segmenting and classifying the individual nuclei, yielding 90 nuclear features (nuHIFs) that quantify shape, size and texture at the single-cell level^39^.

For model development, we analyzed 556 patient samples from TCGA-BRCA with HIFs and nuHIFs, matched with the status of the three clinical markers (HER2, ER and PR) used to define tumor subtype and guide treatment in BC (**Supplementary Tables 1-2**). We applied the widely used Nottingham grading (NHG) criteria^41,42^ (see **METHODS**) to obtain 25 features that are directly interpretable by pathologists, constructing a compact set of *pathologist-interpretable features* (PIFs) (**Supplementary Table 3**). We further focused on the 12 most interpretable PIFs associated with the nuclear morphology, yielding the *nuclear pathologist-interpretable features* (NPIFs; **Supplementary Table 4**). We used these feature sets (PIFs, NPIFs) to build simple, interpretable machine learning (ML) pipelines for prediction (**Figure 1A**).

**Figure 1:**
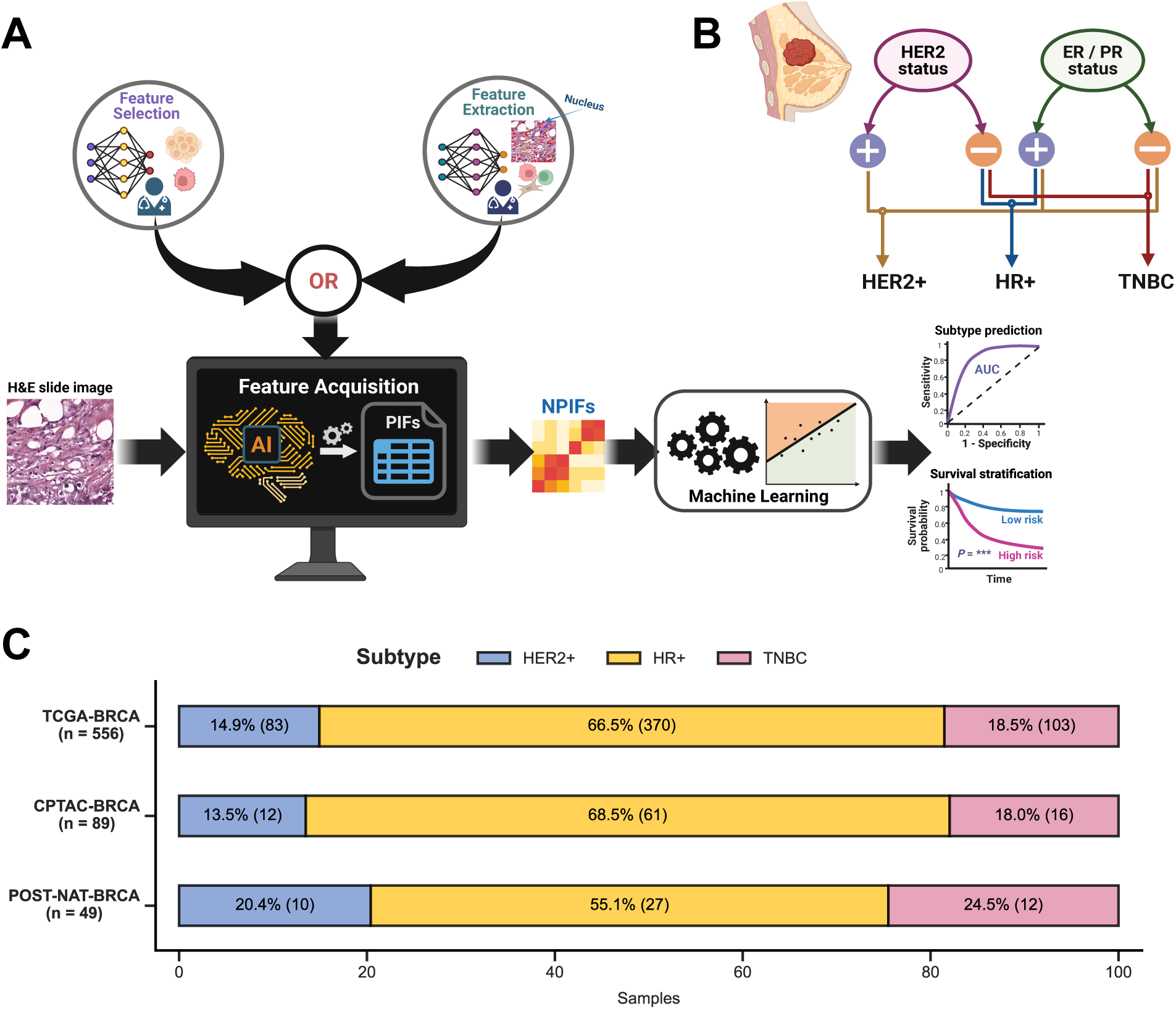
Overview of the methodology: pipeline and cohorts. **A.** Overview of analysis pipeline, EXPAND. *First*, the nuclear pathologist-interpretable features (NPIFs) from hematoxylin and eosin (H&E)-stained whole-slide images are obtained through the feature acquisition module (either two-stage selection from already-extracted HIFs and nuHIFs, or direct extraction from nuclear morphology *via* segmentation); *Second*, we train machine learning pipelines with NPIFs to predict tumor subtypes or patient survival and evaluate performance by using the area under the receiver operating characteristics curve (AUC) metric or Kaplan-Meier analysis and Log-rank test, respectively. **B.** Breast tumor clinical subtype definition based on the statuses of three clinically established markers: HER2 (human epidermal growth factor receptor 2), ER (estrogen receptor) and PR (progesterone receptor). **C.** Distribution of tumor subtypes across the histopathology cohorts used for analysis.

We focused on the three clinically actionable BC subtypes (**Figure 1B**; see **METHODS**): *HER2-positive (HER2+)*, *hormone receptor-positive (HR+)* and *triple-negative breast cancer (TNBC).* For subtype prediction, we devised a transparent ML pipeline to build three dedicated logistic regression (with lasso regularization) models to predict the three subtypes. For survival analysis, we adapted the ML pipeline to build Cox regression (with an optional collinearity removal step before) and Kaplan-Meier models to stratify the subtype-specific survival (**Figure 1A**). The pipeline uses nested cross-validation to optimize model hyperparameters and ensure robustness (see **METHODS**). For state-of-the-art comparison, we train models using HIFs, nuHIFs, and DL-derived direct WSI features, all processed through the same pipeline. To demonstrate model generalizability, we externally validated EXPAND on two completely independent cohorts: CPTAC-BRCA^43^ and POST-NAT-BRCA^44^. Notably, both TCGA-BRCA and CPTAC-BRCA consist of pretreatment samples whereas POST-NAT-BRCA comprises posttreatment specimens following neoadjuvant therapy. Finally, we developed and studied a further fine-grained four subtype classification (*i.e.*, triple-positive breast cancer (TPBC), HER2+, HR+ and TNBC; **Supplementary Figure S1A**; see **METHODS**)^45^.

### Pathologist-interpretable compact EXPAND models identify breast tumor subtypes as accurately as previously published much larger models

We first identified the minimal set of HIFs and nuHIFs^38,39^ that characterize the tumor microenvironment, obtained from H&E-stained WSIs. HIFs quantify different aspects of tissue structure and cell appearance in tumor WSIs such as the tissue region properties, cell concentration, spatial organization and so on. Broadly, they are grouped into seven categories: area, density, density ratio, total count, count proportion, region properties and cell clustering properties (**Supplementary Table 5**). Similarly, nuHIFs capture detailed nuclear morphology across three cell types (cancer cells, fibroblasts and lymphocytes) encompassing the statistical summary (mean and standard deviation; SD) of 15 key characteristics including size, shape, texture and stain intensity (**Supplementary Tables 6**-**7**). These features offer to improve the interpretability of the cellular microenvironment and provide valuable insights into cancer pathology.

Based on the expert opinions from our team of pathologists, we leveraged the established NHG criteria to construct the set of *pathologist-interpretable features* (PIFs). We selected 25 features where each feature reflects one of the three NHG components: nuclear pleomorphism, mitotic count and tubule formation^41,42^ (**Figure 2A, Supplementary Table 8**; see **METHODS**). Furthermore, to maximize model interpretability and generalizability, we focused on nuclear pleomorphism (the most pathologist-interpretable characteristic) to obtain a compact set of 12 *nuclear PIFs* (NPIFs; **Figure 2A, Supplementary Table 9**). To evaluate the informativeness of these compact feature sets, we trained and evaluated the EXPAND subtype classifiers using all four feature sets: First, models based on (1) PIFs (*m* = 25) and (2) NPIFs (*m* = 12), which are the main goal of this study. Second, we built models based on (3) HIFs (*m* = 607) and (4) nuHIFs (*m* = 90), which are trained on the large sets of features already published and serve at control models for comparison. As an additional control model, we compared to DL-based direct WSI-derived features (“direct”, *m* = 512), as extracted by DeepPT, our recently published work^15^.

**Figure 2:**
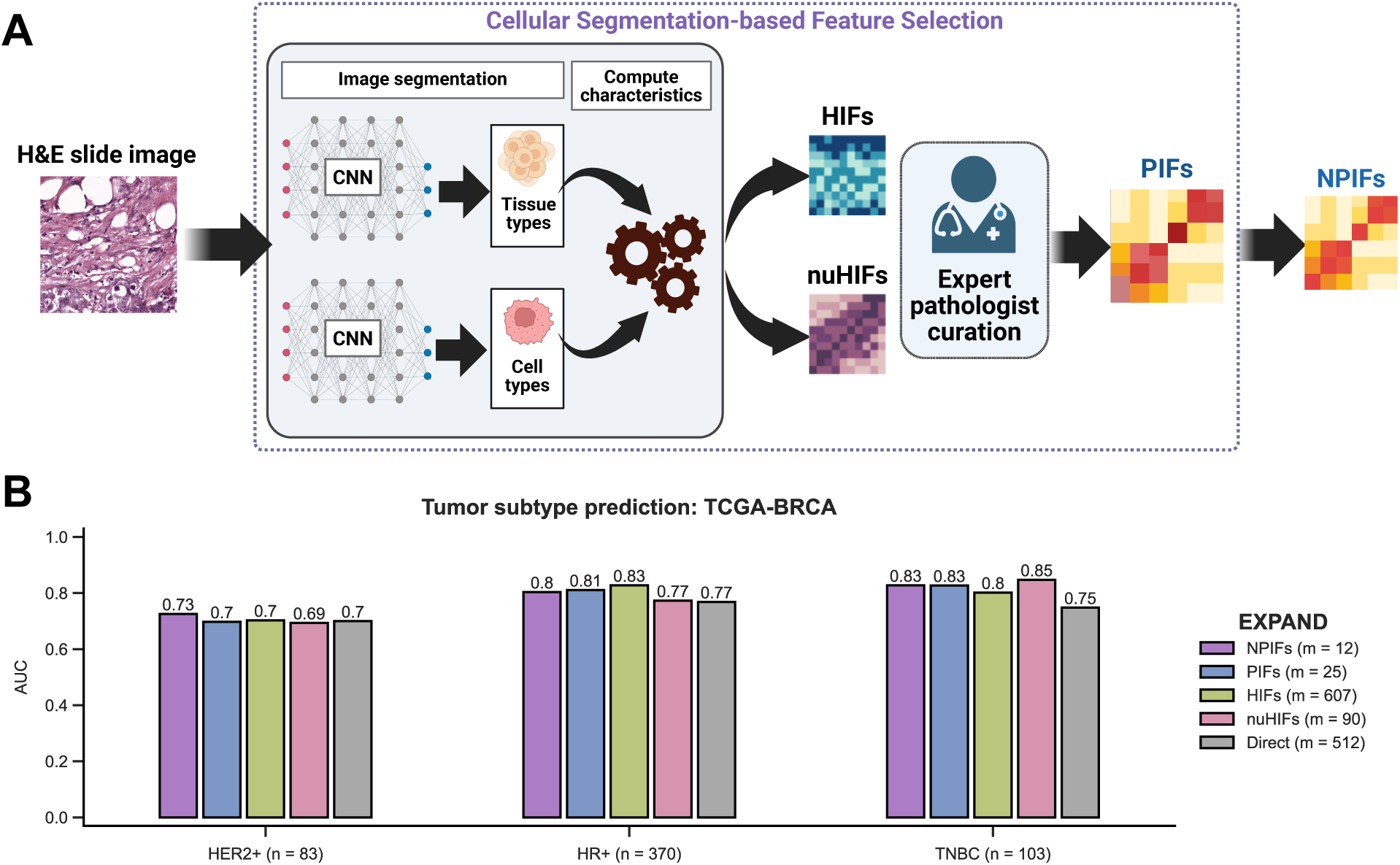
Pathologist-interpretable features (PIFs) and nuclear PIFs (NPIFs) identify tumor subtypes with high accuracy while maintaining interpretability. **A.** Detailed pipeline for the extraction of PIFs and NPIFs based on extracted interpretable features. HIFs and nuHIFs were originally derived using CNN-based frameworks that were trained using H&E-stained WSIs paired with pathologist-annotated cell and tissue types^38,39^. Subsequently, 25 PIFs were curated based on the three NHG criteria and a further filtering yielded a compact set of 12 NPIFs with the highest interpretability. CNN, H&E, WSI and NHG stand for convolutional neural network, hematoxylin and eosin, whole-slide image and Nottingham grading, respectively. **B.** Performance comparison for predicting tumor subtypes in TCGA-BRCA (*n* = 556) using five feature sets: NPIFs, PIFs, HIFs, nuHIFs and direct WSI-derived features. ‘*m*’ denotes the feature set size. AUC stands for the area under the receiver operating characteristics curve.

The EXPAND models based on the compact feature sets perform well. The one based on PIFs obtained area under the receiver operating characteristics curve (AUC) of 0.70, 0.81, 0.83 and those based on NPIFs achieved AUCs of 0.73, 0.80, 0.83 to predict HER2+, HR+ and TNBC tumors, respectively (**Figure 2B**). Reassuringly, when analyzing the control models’ performance, the EXPAND models based on HIFs yielded similar values. The HIF model AUCs were 0.70, 0.83 and 0.80. Similarly, nuHIFs-based EXPAND models achieved comparable AUCs (0.69, 0.77, 0.85; **Figure 2B**). Finally, the baseline direct, non-interpretable features attained the lowest AUCs (0.70, 0.77, 0.75; **Figure 2B**). While the performance for PIFs and NPIFs remain *on par* with the high-dimensional HIF sets, their improved clinical interpretability offers significant advantage in real-world diagnostic settings. We performed similar analyses by considering the more fine-grained four subtypes classification (**Supplementary Figures S1B**), where EXPAND retained its high performance when using PIFs and NPIFs (**Supplementary Figures S1C**).

### EXPAND maintains its predictive performance also as an end-to-end pipeline automatically extracting NPIFs *de novo*

Despite balancing interpretability and accuracy, our compilation of NPIFs from HIFs and nuHIFs employs proprietary PathAI software, making it impossible to apply as is to new datasets, where these features are not publicly available. We hence adapted EXPAND to include an ability to automatically extract the 12 NPIFs directly from the pathology images. To this end, we applied Hover-Net, an open-source nuclear segmentation tool. The latter employs a CNN trained with pathologist-annotated H&E-stained whole-slide images to identify and classify the nuclei present in a given image across five major cell types^46^.

To this end, EXPAND employs a three-step nuclear segmentation and feature extraction pipeline (**Figure 3A**): *First*, each WSI was divided into non-overlapping tiles (512×512 pixels at 20x magnification; see examples in **Figure 4**) and Hover-Net was applied to segment each tile separately; *second*, for each cancer cell nucleus identified, six key morphology characteristics were computed by considering the contour identified including the major and minor axis lengths, perimeter, area, eccentricity and circularity; *third*, the tiles were ranked by their cancer cells nuclei abundance (*i.e.*, representing the tumor-enriched regions) to obtain an optimal subset of tiles to estimate the NPIFs as mean and SD of the above tile-level cancer cells characteristics. We used top 25% cancer cell enriched tiles for further downstream analysis, characterizing the NPIFs of high tumor purity regions in the tumor microenvironment (**Supplementary Figure S2C-D**).

**Figure 3:**
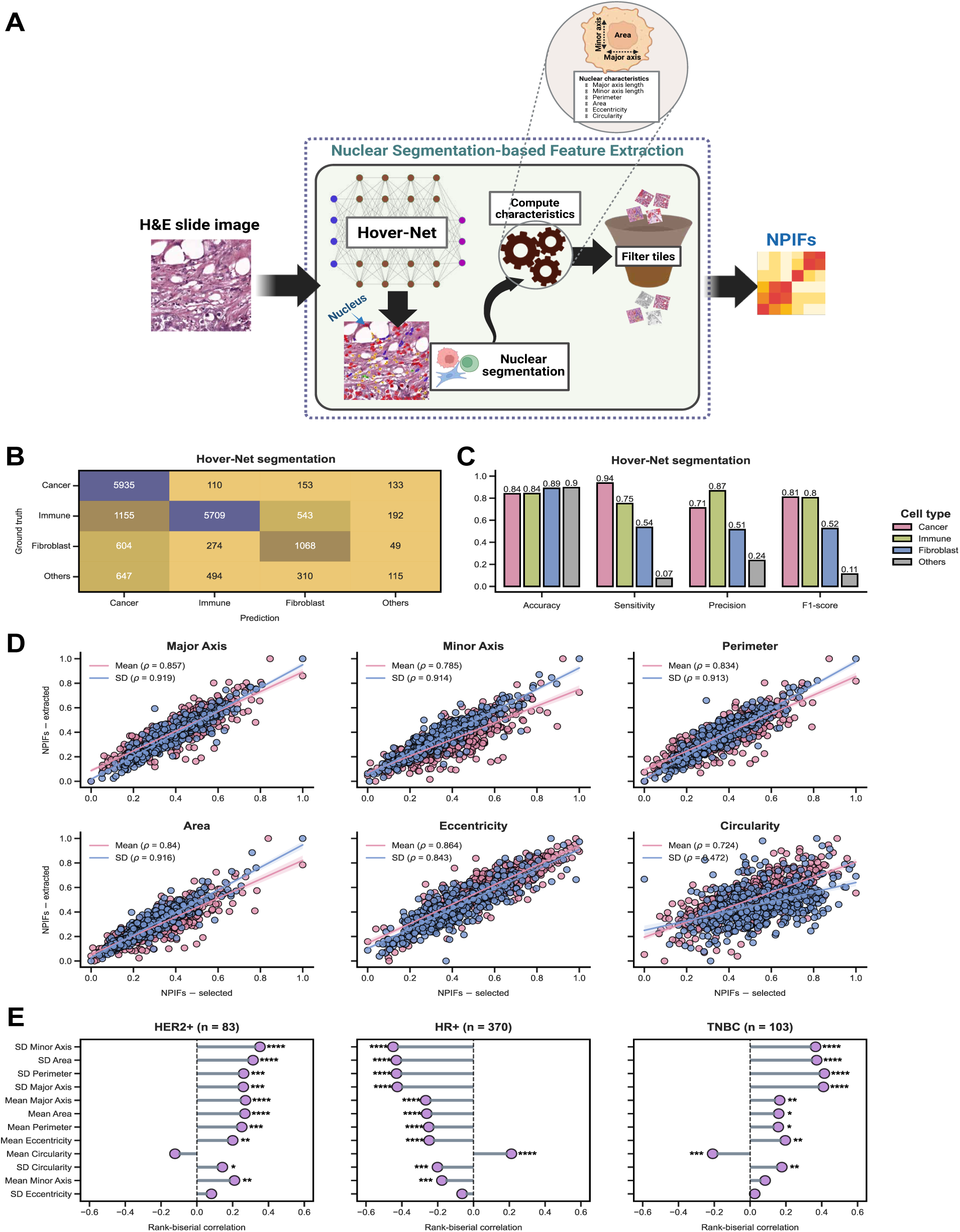
EXPAND facilitates the robust extraction of the NPIFs from whole-slide images. **A.** Detailed pipeline for the segmentation-based extraction of the nuclear pathologist-interpretable features. *First*, each patient WSI is divided into multiple tiles followed by nuclear segmentation using Hover-Net; *Second*, cancer nuclei are isolated to compute six core morphological features; *Third*, for each patient, NPIFs are generated by calculating the mean and SD of these six key morphology characteristics across the top 25% tumor-enriched tiles. H&E and CNN stand for hematoxylin and eosin and convolutional neural network, respectively. **B-C.** Hover-Net nuclear segmentation performance on the PanopTILs dataset (*n* = 151), depicted by confusion matrix (**C**) and per-class performance scores (**D**). Prediction and ground truth denote the Hover-Net identified nucleus types and the pathologist-annotated nucleus types, respectively. Others denote the aggregated class for nuclear types with limited representation (*i.e.*, dead, epithelial and unknown). **D.** Concordance between the EXPAND-selected NPIFs (from HIFs and nuHIFs) and EXPAND-extracted NPIFs for TCGA-BRCA, grouped by the six key morphology characteristics. *ρ* denotes the Spearman correlation coefficient. All values showed are statistically significant (*P* << 0.0001). **E.** Association of the EXPAND-extracted NPIFs with tumor subtypes in TCGA-BRCA, computed by using two-sided Wilcoxon rank-sum tests (****, ***, ** and * denote *P* ≤ 0.0001, *P* ≤ 0.001, *P* ≤ 0.01 and *P* ≤ 0.05), where the effect size is measured by rank-biserial correlation coefficient. NPIFs are ordered by their mean absolute effect sizes across subtypes in a descending order.

**Figure 4.**
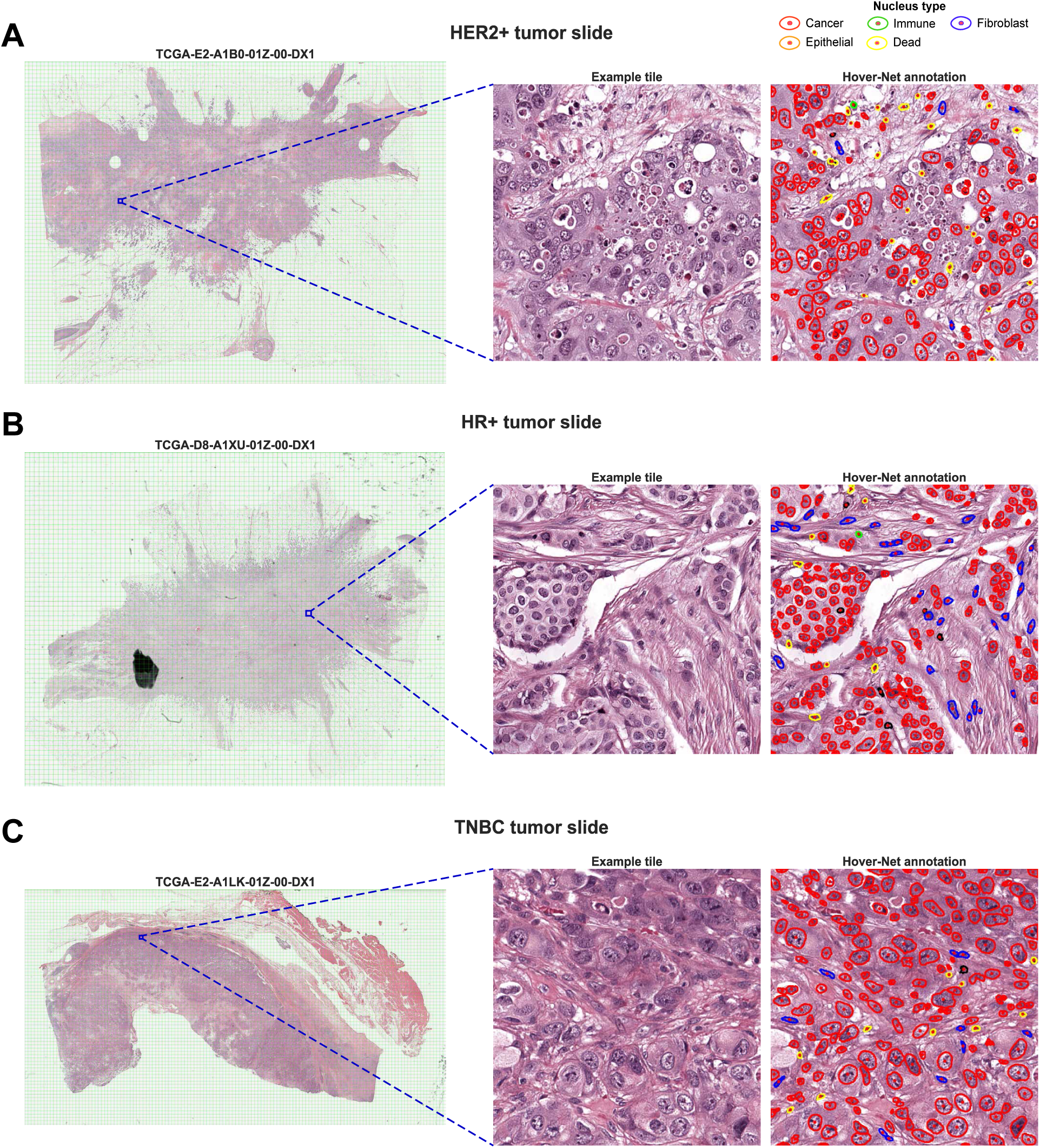
Representative examples of nuclei segmentation using Hover-Net on H&E-stained whole-slide images (WSIs) from TCGA-BRCA. **A.** A representative WSI containing HER2+ breast tumor with a highlighted example tile and the corresponding Hover-Net segmentation results. ***Left***, WSI divided into non-overlapping tiles by the intersecting green lines, where a particular tumor-enriched tile is highlighted as a blue square; ***Middle***, the selected tile displayed at high-resolution; ***Right***, Hover-Net identified cancer cell nuclei (red) within the selected tile where each contour encloses a specific nucleus. **B-C.** Representative WSIs containing HR+ (**B**) and TNBC (**C**) tumors with highlighted example tiles and their corresponding Hover-Net nuclei segmentation results.

We first validated this pipeline by evaluating the nuclear segmentation using PanopTILs, a pathologist-annotated BC dataset (*n* = 151)^40,47^ and then assessing the consistency of the extracted NPIFs with the originally-selected NPIFs. Hover-Net achieved high accuracy and high-to-moderate sensitivity across the three major cell-types nuclei reported in PanopTILs, attesting its robustness (**Figure 3B-C, Supplementary Figures S2A-B**). Further, comparing the extracted NPIFs to the NPIFs previously reported in the TCGA-BRCA achieved near-perfect alignments for five out of six key nuclei features (spearman correlation, *ρ* > 0.8, *P* << 0.0001; **Figure 3D**). Only circularity showed a weaker correlation, especially when computing SD (*ρ* = 0.472, *P* << 0.0001).

To elucidate the biological relevance of the NPIFs, we assessed their association with the tumor subtypes using two-sided Wilcoxon rank-sum tests (*P* ≤ 0.05) and measured the effect size by rank-biserial correlation (**Figure 3E**). We found that both HER2+ and TNBC tumors are positively associated with high levels of variation in nuclear morphology (SD Minor Axis, SD Area, SD Perimeter *etc.*), hallmarks often noted by pathologists when identifying high-grade, aggressive phenotypes (**Figure 4A,C**). In contrast, HR+ tumors have lower levels of these features, consistent with the more uniform and less pleomorphic nuclear shape associated with this subtype (**Figure 4B**). This inverse pattern across subtypes strongly reflects the diagnostic intuition, where the presence of irregular, enlarged and heterogeneously shaped nuclei signals biologically aggressive disease, and the more uniform nuclear architecture typifies well-differentiated, hormone-responsive tumors^48–50^. Similar patterns were observed by considering the fine-grained four subtypes (**Supplementary Figure S2E**).

**Table 1:**
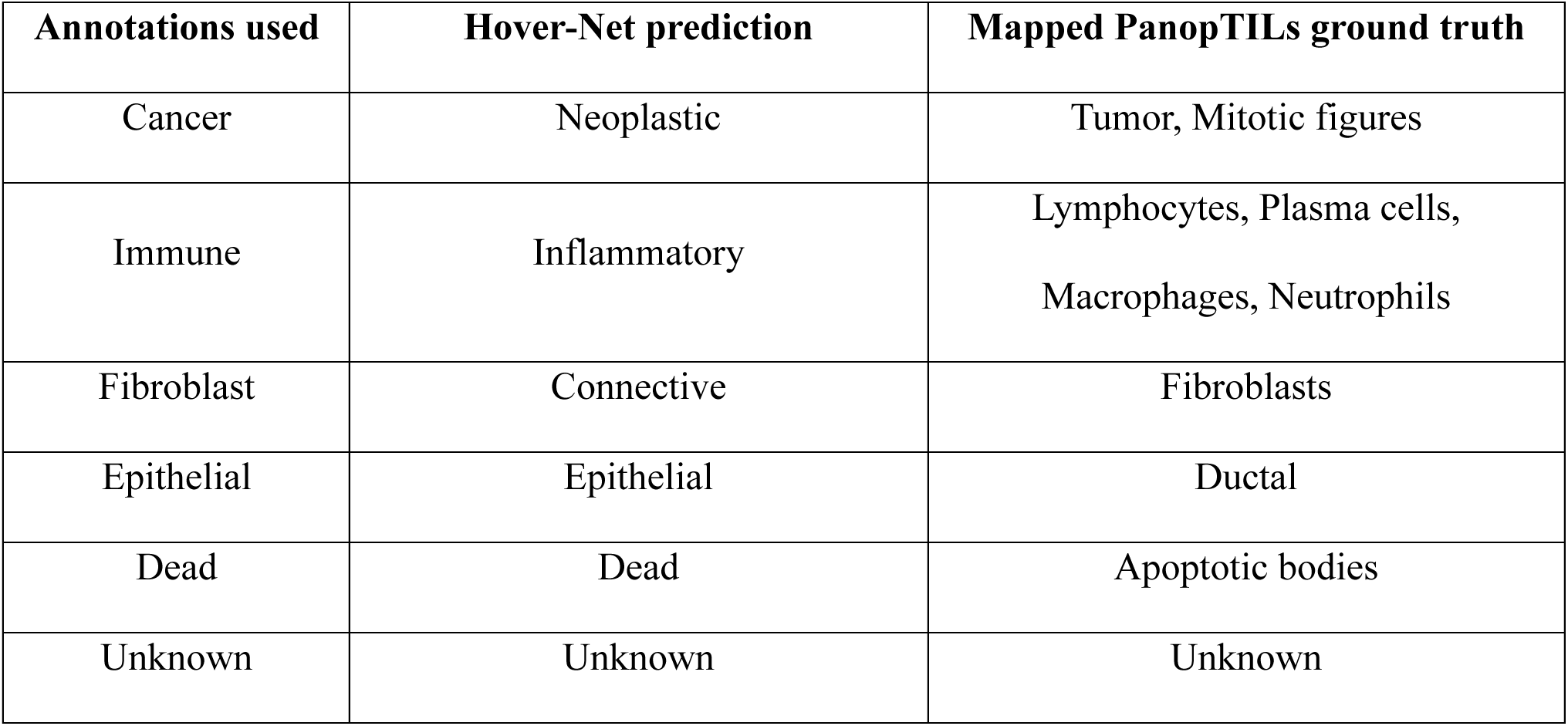
Mapping between Hover-Net predicted cell types and PanopTILs cell types.

### EXPAND reliably maintains subtype prediction accuracy in independent unseen pre and post treatment cohorts

After constructing and validating the automatic segmentation and feature extraction component of EXPAND, we turned to test our ability to use EXPAND-extracted NPIFs to predict tumor subtypes using these features. In the TCGA-BRCA (**Figure 5A, Supplementary Figure S3A**) these extracted NPIFs yielded similar AUCs for both HER2+ and HR+ tumors but slightly lower AUC for TNBC (**Figure 5A**) to those obtained using the previously annotated nuclei features. The decreased performance in TNBC may result from their increased heterogeneity and lower predictive ability of the nuclei features (*e.g.*, SD Circularity (**Figure 3D**). Here again, selecting the top 25% tumor-enriched tiles to compute the NPIFs increased the predictive performance (**Supplementary Figure S2D**).

**Figure 5:**
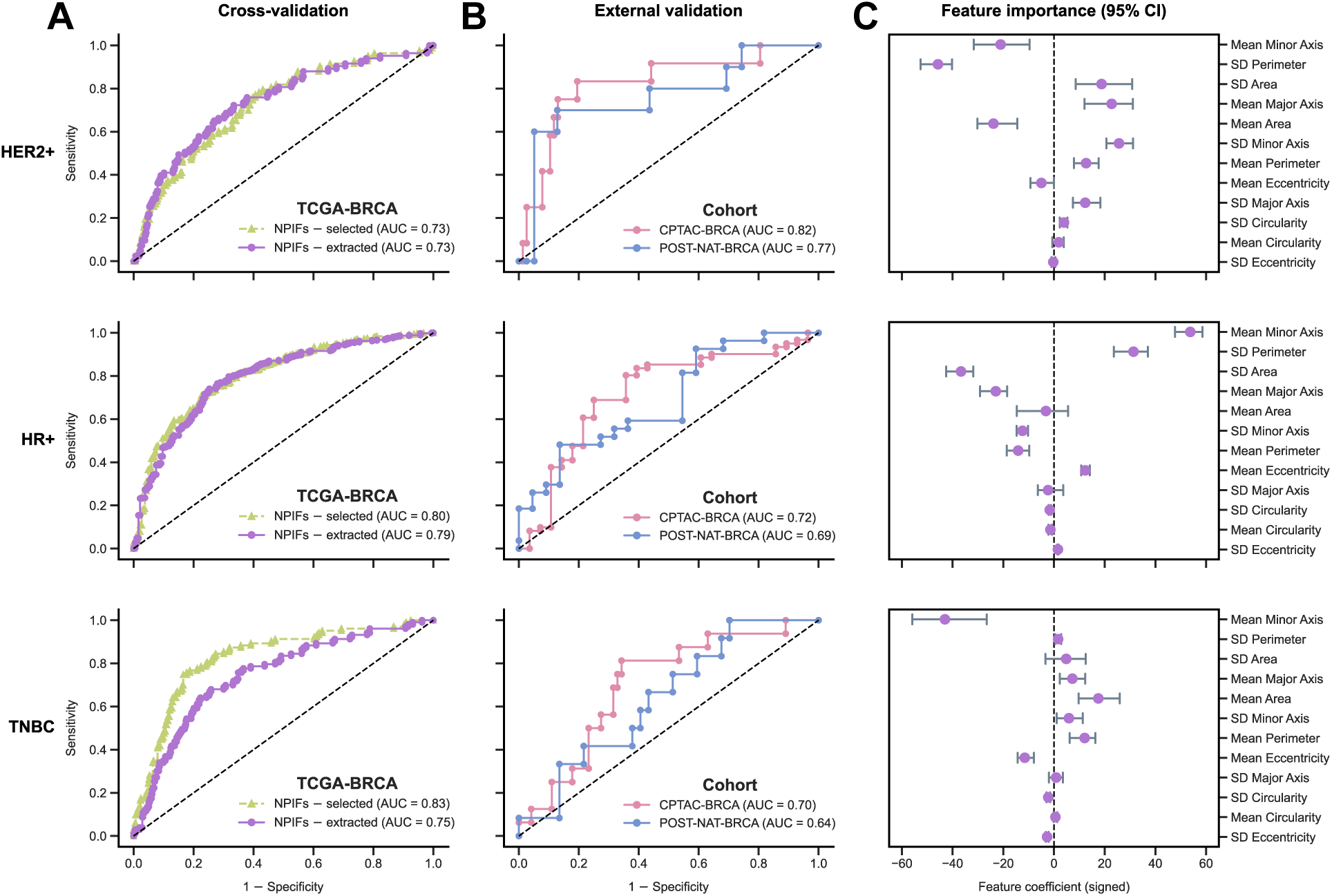
EXPAND reliably identifies tumor subtypes using both selected and extracted NPIFs, and further generalizes to external cohorts. **A**. Tumor subtype prediction in TCGA-BRCA (*n* = 556) using EXPAND with the selected and extracted NPIFs. AUC stands for the area under the receiver operating characteristics curve. **B**. External validation of EXPAND subtype classifiers on the independent cohorts: CPTAC-BRCA (*n* = 89) and POST-NAT-BRCA (*n* = 49). **C**. Feature importances (mean with 95% confidence interval; CI) for the NPIFs in tumor subtype classification, where higher values indicate high impacts on prediction and *vice versa*, with the sign denoting the direction of impact. NPIFs are ordered by their mean absolute feature importance across subtypes in a descending order.

We next assessed the generalizability of EXPAND in external validation in two independent patient cohorts, CPTAC-BRCA (*n* = 89) and POST-NAT-BRCA (*n* = 49). Across both cohorts, the subtype classifiers maintained their performance, achieving AUCs of 0.82, 0.72, 0.70 for CPTAC-BRCA and 0.77, 0.69, 0.64 for POST-NAT-BRCA when predicting HER2+, HR+ and TNBC tumors, respectively (**Figure 5B**). Remarkably, these two cohorts represented two different time-points likely affecting cellular and nuclear morphology. These results thus confirm the robustness and transferability of EXPAND across diverse populations, prevailing small sample sizes and treatment-induced changes. EXPAND further yielded a comparable performance in predicting the fine-grained four subtypes (**Supplementary Figures S3A-B**).

To further elucidated the contribution of the individual segmentation-inferred NPIFs in subtype prediction, we assessed the importance of the NPIFs (mean with 95% confidence interval; CI) across each subtype (**Figure 5C**), showing similar trends to the results shown previously feature association analysis (**Figure 3E**). Both *Mean Area* and *SD Area* showed high importances for HER2+ tumors but only one of these features showed a significantly higher importance for HR+ (*SD Area*) and TNBC (*Mean Area*) tumors, which is congruent with the irregular nuclear shapes observed across HER2+ and TNBC tumors compared to the HR+ tumors. Similar patterns were also observed for the fine-grained four subtypes (**Supplementary Figure S3C**). Overall, these features reflect the intercellular variability in nuclear morphology, characteristics that pathologists routinely associate with tumor phenotypes.

We also explored if EXPAND can be further refined by considering the morphology of the immune nuclei. To this end, we first extracted the 12 “Immune NPIFs” by applying our Hover-Net-based methodology. We then trained and evaluated the EXPAND subtype classifiers on TCGA-BRCA but achieved inferior performance to that obtained with the original cancer-based NPIFs, both when using the top 25% immune-enriched tiles (**Supplementary Figure S4A-B**) or all available tiles (**Supplementary Figure S4C-D**) for NPIF inference. Furthermore, assessing these immune-based classifiers yielded poor generalization for TNBC across both CPTAC-BRCA and POST-NAT-BRCA (**Supplementary Figure S4**). Notably, these results reinforce the longstanding pathology practice of focusing on cancer cells on histopathology slides as the main indicator of patient disease phenotypes.

Collectively, these findings reinforce that EXPAND retains strong predictive performance while offering flexible feature acquisition, high interpretability, accessibility and reproducibility, thus offering a clinically adaptable approach for BC subtype stratification.

### EXPAND successfully stratifies patient survival

We finally turned to study the ability of EXPAND to generate interpretable predictors of BC patient survival by systematically comparing its predictive power vs out control, previously published HIFs and nuHIFs predictors. Since different subtypes are associated with different BC prognosis, we adapted EXPAND to model patient survival for each subtype separately. Briefly, for each subtype, we built a multivariate Cox regression model that used a given feature set (post collinearity filtering) and patient age (as a confounder) to derive the subtype-specific risk score (**Figure 1A**). Subsequently, we stratified patients into Low– and High-risk groups by applying a fixed threshold of 0.5 on the quantile-normalized risk score (using the [10%, 90%] interval; see **METHODS**) and used Kaplan-Meier analysis to demonstrate this stratification for both overall survival (OS; **Figure 6A-C**) and progression-free survival (PFS; **Supplementary Figure S5**).

**Figure 6:**
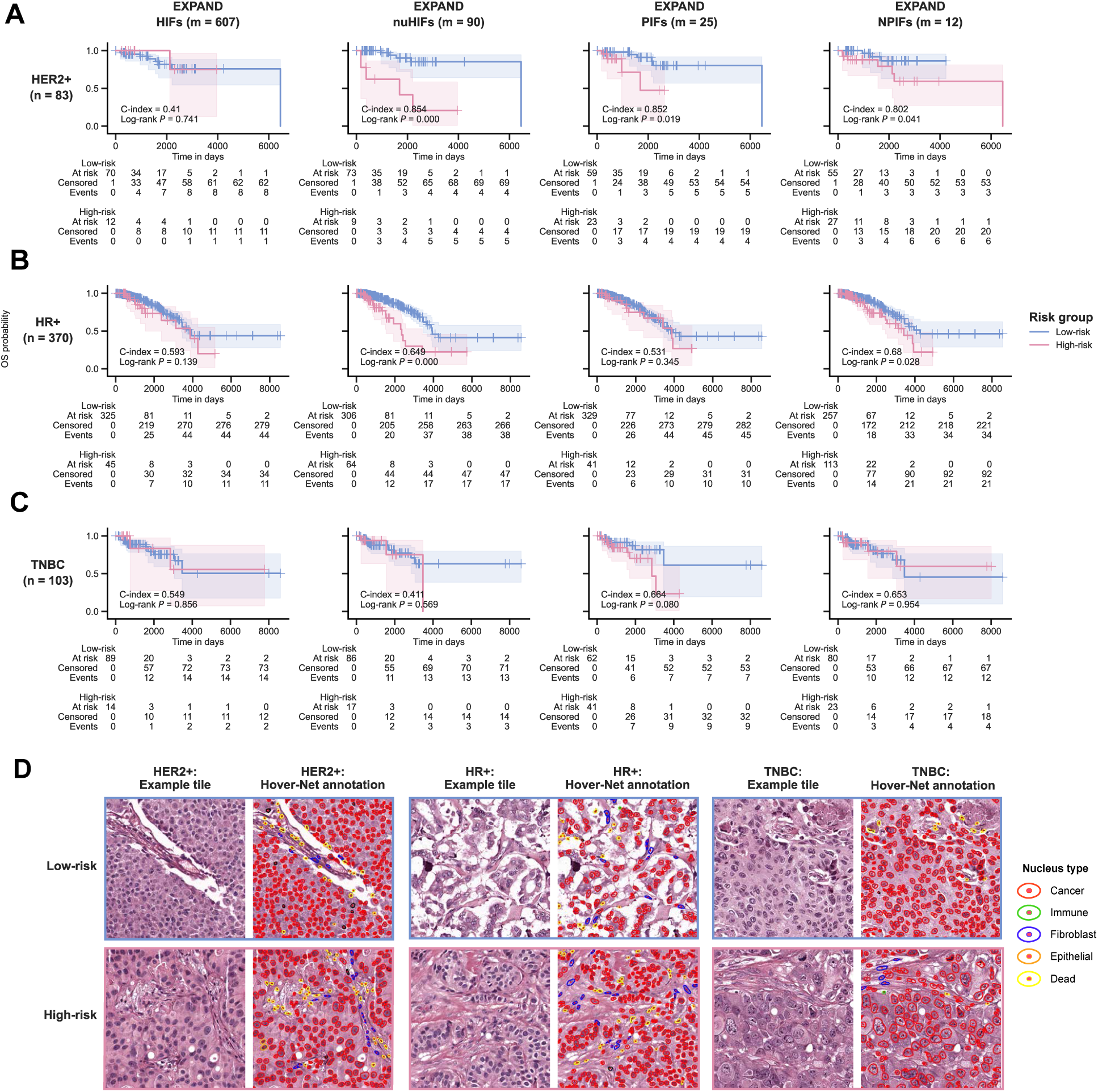
EXPAND stratifies TCGA-BRCA patient survival within each tumor subtype. **A-C.** Kaplan-Meier curves depicting the overall survival (OS) probabilities for TCGA-BRCA patients with HER2+ (**A**), HR+ (**B**) and TNBC (**C**) tumors, stratified using models built with HIFs, nuHIFs, PIFs and NPIFs. Patients were stratified into ‘Low-risk’ and ‘High-risk’ groups by using a fixed threshold of 0.5 on quantile-normalized risk score (using the [10%, 90%] interval). The differences between two curves were computed by using a Log-rank test (*P* ≤ 0.1). **D**. Representative H&E-stained tile images (**left**) and matched Hover-Net annotations (**right**) for TCGA-BRCA patients with low and high predicted survival risks across subtypes.

As displayed in **Figure 6A-B**, EXPAND effectively stratified HER2+ and HR+ patients (*P* ≤ 0.1) using nuHIFs, PIFs and NPIFs with high C-indices (HER2+ = 0.854, 0.852, 0.802; HR+ = 0.649, 0.531, 0.680), while using HIFs failed to significantly stratify across either subtype (HER2+ = 0.410, HR+ = 0.593). Notably, EXPAND faced a significant hurdle to stratify TNBC patients using HIFs (0.549), nuHIFs (0.411) and NPIFs (0.653) but achieved a significant fit for PIFs (0.664, *P* ≤ 0.1; **Figure 6C**), suggesting that the texture and stain intensity information, alongside the nuclear features, could be necessary to predict TNBC prognosis. **Figure 6D** shows some representative H&E-stained tile images and their matched nuclear annotations across risk groups for each subtype. Similar survival trends were observed when stratifying for PFS (**Supplementary Figure S5**). These results highlight the prognostic capabilities of the PIFs and NPIFs compared to the larger HIF sets, whereas the need for additional features beyond nuclear characteristics for TNBC underscores the increased challenge inherent to this subtype.

## DISCUSSION

Our study introduced EXPAND, a first of its kind compact and interpretable end-to-end approach to identify the breast tumor subtype and prognosis from a patient’s histopathology image, that is based on features that are commonly used by pathologists in the routine workflow. EXPAND was designed to obtain a small set of *pathologist-interpretable features* (*PIFs*) that encode morphological characteristics routinely assessed by pathologists, thus balancing model interpretability with high performance. EXPAND emphasizes the *nuclear PIFs (NPIFs)*, the most compact and interpretable features that achieved high accuracy and can be extracted using open-source segmentation tools, offering a readily available scalable solution that will be made available for public use upon publication.

We trained EXPAND on TCGA-BRCA to identify the three clinically actionable subtypes (HER2+, HR+ and TNBC) and achieved performance *on par* with proprietary models based on much larger and complex sets of interpretable features, including HIFs and nuHIFs. When applied *as-is*, EXPAND successfully identified tumor subtypes in CPTAC-BRCA and POST-NAT-BRCA external cohorts, demonstrating generalization without overfitting. Remarkably, EXPAND achieved this generalization across both pretreatment (CPTAC-BRCA) and posttreatment (POST-NAT-BRCA) samples, thus overcoming the treatment-induced changes likely present in POST-NAT-BRCA, despite only being trained on pretreatment TCGA-BRCA samples. Furthermore, the associations of NPIFs with different subtypes match common knowledge by pathologists, where HER2+ and TNBC tumors are known to exhibit more nuclear pleomorphism whereas HR+ tumors are characterized by more uniform and well-differentiated nuclei^48–50^. Notably, our use of the top 25% tumor-enriched tiles within a WSI for computing NPIFs also concords with the typical pathology workflow, where expert pathologists would focus on the diagnostically relevant areas during slide review^51^.

While we demonstrated the robustness and generalizability of PIFs and NPIFs, certain limitations should be acknowledged and addressed in future studies. *First*, EXPAND is built using primary tumor WSIs, potentially limiting generalizability to metastatic tumors. This limitation is inherent to DL-based feature acquisition, as most segmentation tools including Hover-Net are developed using primary WSIs and exhibit suboptimal performance on metastatic WSIs^52^. *Second*, EXPAND is designed to align with the routine pathology evaluation by focusing on the cancer nuclei morphology without explicitly accounting for the spatial interactions among diverse cell types within the tumor microenvironment (TME). This likely constrains EXPAND’s ability to characterize intra-tumor heterogeneity closely, possibly limiting the performance observed in histologically complex subtypes like TNBC^53,54^.

Building upon the success of EXPAND in breast cancer, one future direction would be to extend the use of the NPIFs to other cancers, capturing morphological differences across tumor types. This could potentially include additional major cell types in the TME (*e.g.,* immune cells, stromal cells *etc.*), to capture more complex dynamics. Another intriguing direction would be to build a high-resolution pathologist-interpretable AI model by leveraging tile-level characteristics coupled with attention-based algorithms, integrating both top-down and bottom-up approaches.

Finally, integrating EXPAND with multiomics data could potentially guide towards interpretability-aware multimodal modeling that better reflects the complex interplay between tumor morphology and molecular phenotypes, thus improving precision oncology applications.

In summary, EXPAND presents a notable advance in integrating computational pathology with clinical diagnostics by introducing a transparent, interpretable end-to-end AI framework for breast tumor subtyping and risk stratification. Designed to focus on nuclear morphology, a key component of pathology diagnosis, these NPIFs offer a reliable and scalable way to aid in routine pathological assessment, potentially accelerating it. The relative contributions of different features in prediction could further assist pathologists focus on areas of interest, thus potentially enriching the knowledge base for humans and AI alike. The successful generalization of EXPAND to external cohorts highlights its translational value and supports its further prospective testing for clinical integration. This approach could serve as a guide for developing a pan-cancer pathologist-interpretable model, making AI in pathology more transparent, practical and aligned with real-world clinical decision-making.

## METHODS

### Data overview

This study utilized H&E-stained whole-slide images and corresponding image-derived feature sets from three publicly available breast cancer cohorts: The Cancer Genome Atlas (TCGA-BRCA) for model development, Clinical Proteomic Tumor Analysis Consortium (CPTAC-BRCA)^43^ and post neoadjuvant treatment of breast cancer (POST-NAT-BRCA)^44^ for external validation.

**TCGA-BRCA:** 556 patients were available with both human-interpretable image features, derived from their WSIs (*n* = 556), and matched HER2, ER and PR statuses (positive / negative). Using these statuses, the patient tumors were classified into three clinically actionable subtypes (**Figure 1B**):

- **HER2-positive (HER2+):** tumors with HER2-positive status.
- **Hormone receptor-positive (HR+):** tumors with HER2-negative and ER-positive or PR-positive statuses.
- **Triple-negative breast cancer (TNBC):** tumors with negative HER2, ER and PR statuses.

To further refine tumor subtyping, we applied a fine-grained four subtype classification using the HER2 and ER statuses alone^45^. ER-positive and ER-negative were treated as ER+/PR+ and ER-/PR-, respectively, given their strong correlation (**Supplementary Figure S1A**):

- **Triple-positive breast cancer (TPBC):** ER-positive, HER2-positive tumors
- **HER2-positive (HER2+):** ER-negative, HER2-positive tumors
- **Hormone-receptor positive (HR+):** ER-positive, HER2 negative tumors
- **Triple-negative breast cancer (TNBC):** ER-negative, HER2-negative tumors

For the three-subtype classification, TCGA-BRCA included 83 HER2+, 370 HR+ and 103 TNBC cases (**Figure 1C**), whereas for the four-subtype classification, 58 TPBC, 25 HER2+, 361 HR+ and 112 TNBC cases were identified (**Supplementary Figure S1B**).

**CPTAC-BRCA:** 89 patients were available with matched WSIs (*n* = 168) and clinical marker statuses^43^. Among them, 12 HER2+, 61 HR+ and 16 TNBC cases were identified for the three-subtype classification (**Figure 1C**), while the four-subtype classification comprised 7 TPBC, 5 HER2+, 59 HR+ and 18 TNBC cases (**Supplementary Figure S1B**). For patients with multiple WSIs (varying from 1 to 4), the multiple WSI-level features were averaged to obtain the patient-level features.

**POST-NAT-BRCA:** 49 patients were available with matched WSIs (*n* = 94), collected following neoadjuvant therapy, and clinical marker statuses^44^. Among them, 10 HER2+, 27 HR+ and 12 TNBC cases were identified for the three-subtype classification (**Figure 1C**), while the four-subtype classification comprised 8 TPBC, 2 HER2+, 27 HR+ and 12 TNBC cases (**Supplementary Figure S1B**). For patients with multiple WSIs (varying from 1 to 2), the multiple WSI-level features were averaged to obtain the patient-level features.

### Feature acquisition

We investigated three classes of interpretable features that were obtained from patient WSIs to build our predictive models in downstream analyses:

- **HIFs:** 607 human-interpretable image features that capture the tissue– and cell-level characteristics, developed by Diao et al^38^.
- **nuHIFs:** 90 nuclear human interpretable features that quantify the nuclear morphology, developed by Abel et al^39^.
- **PIFs and NPIFs:** 25 pathologist-interpretable features that follow the Nottingham Grading (NHG) criteria and includes a core set of 12 nuclear pleomorphism-based PIFs, both developed in this work.

The HIFs and nuHIFs for TCGA-BRCA were obtained from the corresponding publications^38,39^. While these derived features are publicly available, the original cell– and tissue-level annotations for model training remain proprietary and inaccessible. We also built a baseline control model with 512 deep learning-derived non-interpretable features generated by DeepPT and DEPLOY, our recently published AI frameworks^15,19^. These features were initially extracted by using a ResNet50 pre-trained model^55^ (yielding 2,048 features) and subsequently compressed into 512 compact features with an autoencoder.

We curated the PIFs as features aligned with the three standard criteria in NHG (*i.e.*, nuclear pleomorphism, mitotic count and tubule formation) after consulting with a panel of expert pathologists. Specifically, 12 features reflecting the nuclear size, shape and irregularity were selected by full consensus and referred as the NPIFs.

**NPIFs extraction *via* segmentation:** To enable a scalable feature extraction, we devised an alternative pipeline to obtain the NPIFs using Hover-Net, an open-source nuclear segmentation tool^46^. Each WSI was first divided into non-overlapping tiles of size 512 × 512 pixels (at 20x magnification) and processed according to the image processing and stain normalization protocols outlined in DeepPT^15^ and DEPLOY^19^. Hover-Net was used to segment and classify the nuclei present in each tile. After filtering out the tiles with no cancer nuclei, each nucleus within a tile was treated as an individual instance to compute six key morphological characteristics (*i.e.*, major axis length, minor axis length, perimeter, area, eccentricity and circularity) based on their contour by using the Shapely Python package^56^. The 12 NPIFs were extracted for each WSI by computing the mean and standard deviation (SD) of these six characteristics by considering the cancer nuclei within the top 25% tumor-enriched tiles (*i.e.*, most informative region) for that specific WSI.

To determine the optimal cut-off for the top tumor-enriched tiles for an WSI, we leveraged the already-selected NPIFs (from HIFs and nuHIFs) for TCGA-BRCA as ground truth labels to evaluate the segmentation-based NPIFs. All tiles within a WSI were ranked by their proportion of cancer nuclei (*i.e.*, levels of tumor enrichment) to systematically compare the NPIFs generated from using the top 5% to 50% of the tiles (continuous increase of 5%) and all viable tiles in terms of their correlation with the already-selected NPIFs and modeling performance, with the cut-off of 25% tiles showing the most consistent scores (**Supplementary Figure S2C-D**). We used this cut-off for further analyses involving external cohorts.

### Nuclear segmentation with Hover-Net

We adopted Hover-Net for automatic nuclei segmentation and classification from H&E-stained WSIs, identifying the nuclei for five distinct cell/tissue types: cancer, immune, fibroblast, epithelial and dead (**Figure 3A**). To rigorously assess the segmentation accuracy, we used the PanopTILs dataset, a publicly available dataset containing 815 regions of interest (ROIs) from 151 TCGA-BRCA WSIs, with high-quality ground truth annotations verified by a panel of expert pathologists. These annotations included both nuclear types and their contours, providing a robust benchmark for evaluating Hover-Net performance.

We applied Hover-Net to predict nucleus types and contours across all 815 ROIs in PanopTILs, generating annotations directly comparable to ground truth. The predictions were matched to ground truth based on their contour similarity and nucleus type. Specifically, **f**or each nucleus in PanopTILs, we iteratively searched for the corresponding Hover-Net-predicted nucleus based on both contour similarity and nucleus type. A match was confirmed if the predicted nucleus had an intersection over union (IoU) > 0.5 with a ground truth nucleus of the same type.

To evaluate model performance, we generated a multiclass confusion matrix by first aligning the Hover-Net-predicted nucleus types with the PanopTILs ground truth. Hover-Net generated five distinct annotations: neoplastic, inflammatory, connective, epithelial and dead, whereas ground truth labels included tumor / mitotic figures, lymphocytes / plasma cells / macrophages / neutrophils, fibroblast, ductal and apoptotic bodies. Therefore, to align predictions with ground truth, we devised a map grouping similar categories together, as follows:

Following this, we calculated the standard classification metrics *i.e.*, accuracy, sensitivity, precision and F1-score for each nucleus type (**Figure 3C**, **Supplementary Figure S2B**).

### Machine learning modeling

#### Subtype classification

We developed interpretable, subtype-specific classifiers using logistic regression (LR) with L1 (lasso) regularization, training a separate classifier for each BC subtype using a one-vs-all scheme. This included designing both three-subtype (HER2+, HR+, TNBC) and fine-grained four-subtype (TPBC, HER2+, HR+, TNBC) frameworks. For each classification task, the target subtype was binarized (*e.g.*, 1 if the sample contains a tumor of that subtype and 0 if not) and the classifier was trained using a 5×5 nested cross-validation (CV) to ensure robust performance and prevent data leakage. The final prediction for a given WSI was taken as the subtype with the maximum predicted score. Each feature set (HIFs, nuHIFs, PIFs, NPIFs and direct) was evaluated individually using this standardized classification pipeline.

**Nested cross-validation**: Model development was performed using a 5×5 nested CV, where the inner folds were used for hyperparameter tuning and the outer folds were reserved for evaluation. Specifically, we designed a two-step ML pipeline as follows:

- Within each outer CV fold, the features were scaled in [0, 1] using the min-max scaler and then a grid search was performed using the inner CV folds to determine the optimal regularization parameter (C).
- The classifier was then retrained using the optimal C on the training data in the outer fold to make predictions on the held out test data.

For each subtype, we trained five LR classifiers across the five outer CV folds that were frozen and saved to apply *as-is* on external data. To evaluate CV performance on TCGA-BRCA, we first concatenated predictions from the outer folds to build the complete prediction score vector for each subtype and compared with ground truth labels to calculate the subtype-specific AUCs.

**External validation:** The NPIFs for each external cohort were fed to the trained classifiers and their prediction scores were averaged to compute the final subtype-specific prediction scores, which were then used to evaluate the generalization performance.

**Feature Importance:** The contribution of each individual NPIF to predicting a specific subtype was quantified using the feature coefficients from the trained LR classifiers. We recorded the exact feature coefficients learned across five outer CV folds in training *i.e.*, five coefficients per NPIF. For each NPIF, these coefficients were then used to compute the overall importance as the mean and the spread of the importance as the 95% confidence interval (CI). Further, the sign of these coefficients represented the direction of impact.

#### Survival Analysis

We performed multivariate survival analysis using Cox proportional hazard (CPH) models for each subtype using all interpretable feature sets: HIFs, nuHIFs, PIFs and NPIFs. For HIFs, nuHIFs and PIFs, we observed high degrees of feature multicollinearity causing convergence issues with CPH, and therefore, applied collinearity filtering by removing the most correlated features based on optimized thresholds specific to each feature set. Each model then incorporated the remaining non-redundant features and patient age, a well-established clinical confounder. We used a fivefold CV to train and evaluate the EXPAND survival models for each subtype using each feature set, training models on 80% of the data and testing on the remaining 20%, while ensuring that each test fold contains at least one observed event (*i.e.*, death).

We modeled both overall survival (OS) and progression-free survival (PFS, synonymous with progression-free interval) endpoints using this pipeline. For each endpoint, the risk scores from the test folds were concatenated to generate the complete risk score vector across all patients. To stratify patients by using a fixed classification threshold, we quantile normalized this risk score (*r*) between 0 and 1 within the [10%, 90%] interval (*i.e.*, rescaling between the 10^th^ and 90^th^ percentiles) to remove outlier effects, as shown below:

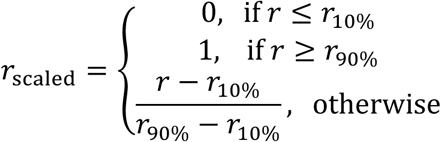

Patients were stratified into ‘Low-risk’ and ‘High-risk’ groups using a fixed threshold of 0.5 on this normalized risk score and their survival differences were depicted using the Kaplan– Meier analysis, where the difference between the two curves were assessed using a Log-rank test (*P* ≤ 0.1).

#### Statistical analysis

To compute the concordance between the selected and extracted NPIFs, we used the Spearman correlation (*P* ≤ 0.05). To compute the association between NPIFs and subtypes, we used the two-sided Wilcoxon test (*P* ≤ 0.05) with the effect size measured by the rank-biserial correlation, computed using the *U*-statistics. To assess the binary classification performance, we analyzed the receiver operating characteristics curve with the effect size measured by the area under the curve (AUC) value. To assess the multiclass classification performance, we computed the multiclass confusion matrix and one vs. all evaluation metrics: Accuracy, Sensitivity, Precision and F1-score. For survival analysis, we performed the Kaplan-Meier (KM) analysis with the risk groups stratified by using a fixed threshold of 0.5 on the quantile-normalized risk score (within the [10%, 90%] interval). The raw risk scores were generated using the multivariate Cox proportional hazard model and used to compute the concordance index (C-index) as the effect size measure. The difference between two KM curves was computed by using the Log-rank test (*P* ≤ 0.1).

#### Implementation details

All analyses were conducted using Python 3.10. Core data preprocessing, machine learning and data visualization were performed using NumPy (v1.24.4), Pandas (v2.0.3), Scikit-learn (v1.3.0), Joblib (v1.3.0), Pickle (v4.0), Matplotlib (v3.7.2) and Seaborn (v0.13.2). Image preprocessing steps (*i.e.*, tile partitioning and color normalization) were performed using OpenSlide (v1.3.1), OpenCV (v4.10.0) and Pillow (v9.2.0). Deep learning models employed PyTorch (v1.12.1). Survival analyses were performed using Lifelines (v0.28.0).

## CODE AVAILABILITY

The EXPAND codes are publicly available at https://github.com/ruppinlab/EXPAND.

## ACKNOWLEDGEMENTS

This research was supported [in part] by the Intramural Research Program of the National Institutes of Health (NIH), National Cancer Institute, Center for Cancer Research. The contributions of the NIH author(s) were made as part of their official duties as NIH federal employees, are in compliance with agency policy requirements, and are considered works of the United States Government. However, the findings and conclusions presented in this paper are those of the author(s) and do not necessarily reflect the views of the NIH or the U.S. Department of Health and Human Services. This work has utilized the computational resources of the NIH HPC Biowulf cluster (http://hpc.nih.gov). The results presented are in part based on the data generated by The Cancer Genome Atlas Research network (https://www.cancer.gov/tcga/). Multiple figure panels (panels A, B in Figure 1, panel A in Figure 2, panel A in Figure 3, panel A in Supplementary Figure S1) have been created with http://biorender.com/. ChatGPT v4o (https://chatgpt.com/) was used strictly to refine certain existing paragraphs for better overall presentation.

## AUTHOR CONTRIBUTIONS

R.K.B. and S.R.D. developed the framework and conducted all analyses. D-T.H., E.D.S., E.M.C., A.T.W., S.A.H., T.C.H., A.P.S., M.P.N. and K.D.A. contributed to the interpretation of results and provided critical feedback throughout the study. E.R. supervised the project. R.K.B., S.R.D. and R. wrote the manuscript with input from all coauthors.

## COMPETING INTERESTS

E.R. is a co-founder of Medaware, Metabomed and Pangea Biomed (divested from the latter). E.R. serves as a non-paid scientific consultant to Pangea Biomed under a collaboration agreement between Pangea Biomed and the NCI. E.R. also serves as a scientific advisory board member of GSK oncology. The other authors declare no competing interests.

## REFERENCES

1 Harbeck, N. et al. Breast cancer. Nat Rev Dis Primers 5, 66 (2019). 10.1038/s41572-019-0111-2

2 Rakha, E. A., Tse, G. M. & Quinn, C. M. An update on the pathological classification of breast cancer. Histopathology 82, 5–16 (2023). 10.1111/his.14786

3 Dunn, C. et al. Quantitative assessment of H&E staining for pathology: development and clinical evaluation of a novel system. Diagn Pathol 19, 42 (2024). 10.1186/s13000-024-01461-w

4 van den Tweel, J. G. & Taylor, C. R. A brief history of pathology: Preface to a forthcoming series that highlights milestones in the evolution of pathology as a discipline. Virchows Arch 457, 3–10 (2010). 10.1007/s00428-010-0934-4

5 Baxi, V., Edwards, R., Montalto, M. & Saha, S. Digital pathology and artificial intelligence in translational medicine and clinical practice. Mod Pathol 35, 23–32 (2022). 10.1038/s41379-021-00919-2

6 Campanella, G. et al. Clinical-grade computational pathology using weakly supervised deep learning on whole slide images. Nat Med 25, 1301–1309 (2019). 10.1038/s41591-019-0508-1

7 Esteva, A. et al. A guide to deep learning in healthcare. Nat Med 25, 24–29 (2019). 10.1038/s41591-018-0316-z

8 Lu, M. Y. et al. AI-based pathology predicts origins for cancers of unknown primary. Nature 594, 106–110 (2021). 10.1038/s41586-021-03512-4

9 Lu, M. Y. et al. Data-èicient and weakly supervised computational pathology on whole-slide images. Nat Biomed Eng 5, 555–570 (2021). 10.1038/s41551-020-00682-w

10 Niazi, M. K. K., Parwani, A. V. & Gurcan, M. N. Digital pathology and artificial intelligence. Lancet Oncol 20, e253–e261 (2019). 10.1016/S1470-2045(19)30154-8

11 Echle, A. et al. Deep learning in cancer pathology: a new generation of clinical biomarkers. Br J Cancer 124, 686–696 (2021). 10.1038/s41416-020-01122-x

12 Al-Tashi, Q. et al. Machine Learning Models for the Identification of Prognostic and Predictive Cancer Biomarkers: A Systematic Review. Int J Mol Sci 24 (2023). 10.3390/ijms24097781

13 Fu, Y. et al. Pan-cancer computational histopathology reveals mutations, tumor composition and prognosis. Nat Cancer 1, 800–810 (2020). 10.1038/s43018-020-0085-8

14 Saltz, J. et al. Spatial Organization and Molecular Correlation of Tumor-Infiltrating Lymphocytes Using Deep Learning on Pathology Images. Cell Rep 23, 181–193 e187 (2018). 10.1016/j.celrep.2018.03.086

15 Hoang, D. T. et al. A deep-learning framework to predict cancer treatment response from histopathology images through imputed transcriptomics. Nat Cancer 5, 1305–1317 (2024). 10.1038/s43018-024-00793-2

16 Johannet, P. et al. Using Machine Learning Algorithms to Predict Immunotherapy Response in Patients with Advanced Melanoma. Clin Cancer Res 27, 131–140 (2021). 10.1158/1078-0432.CCR-20-2415

17 Wang, X. et al. Spatial interplay patterns of cancer nuclei and tumor-infiltrating lymphocytes (TILs) predict clinical benefit for immune checkpoint inhibitors. Sci Adv 8, eabn3966 (2022). 10.1126/sciadv.abn3966

18 Zhang, Y. et al. Histopathology images-based deep learning prediction of prognosis and therapeutic response in small cell lung cancer. NPJ Digit Med 7, 15 (2024). 10.1038/s41746-024-01003-0

19 Hoang, D. T. et al. Prediction of DNA methylation-based tumor types from histopathology in central nervous system tumors with deep learning. Nat Med 30, 1952–1961 (2024). 10.1038/s41591-024-02995-8

20 Arslan, S. et al. A systematic pan-cancer study on deep learning-based prediction of multi-omic biomarkers from routine pathology images. Commun Med (Lond) 4, 48 (2024). 10.1038/s43856-024-00471-5

21 Sammut, S. J. et al. Multi-omic machine learning predictor of breast cancer therapy response. Nature 601, 623–629 (2022). 10.1038/s41586-021-04278-5

22 Li, Y., Wu, X., Fang, D. & Luo, Y. Informing immunotherapy with multi-omics driven machine learning. NPJ Digit Med 7, 67 (2024). 10.1038/s41746-024-01043-6

23 Hollon, T. C. et al. Near real-time intraoperative brain tumor diagnosis using stimulated Raman histology and deep neural networks. Nat Med 26, 52–58 (2020). 10.1038/s41591-019-0715-9

24 Strom, P. et al. Artificial intelligence for diagnosis and grading of prostate cancer in biopsies: a population-based, diagnostic study. Lancet Oncol 21, 222–232 (2020). 10.1016/S1470-2045(19)30738-7

25 Kather, J. N. et al. Deep learning can predict microsatellite instability directly from histology in gastrointestinal cancer. Nat Med 25, 1054–1056 (2019). 10.1038/s41591-019-0462-y

26 Madabhushi, A. & Lee, G. Image analysis and machine learning in digital pathology: Challenges and opportunities. Med Image Anal 33, 170–175 (2016). 10.1016/j.media.2016.06.037

27 Castro, D. C., Walker, I. & Glocker, B. Causality matters in medical imaging. Nat Commun 11, 3673 (2020). 10.1038/s41467-020-17478-w

28 Plass, M. et al. Explainability and causability in digital pathology. J Pathol Clin Res 9, 251–260 (2023). 10.1002/cjp2.322

29 Tjoa, E. & Guan, C. A Survey on Explainable Artificial Intelligence (XAI): Toward Medical XAI. IEEE Trans Neural Netw Learn Syst 32, 4793–4813 (2021). 10.1109/TNNLS.2020.3027314

30 Ali, S. et al. The enlightening role of explainable artificial intelligence in medical & healthcare domains: A systematic literature review. Comput Biol Med 166, 107555 (2023). 10.1016/j.compbiomed.2023.107555

31 Finlayson, S. G. et al. Adversarial attacks on medical machine learning. Science 363, 1287–1289 (2019). 10.1126/science.aaw4399

32 Zhao, Y. et al. Deep learning using histological images for gene mutation prediction in lung cancer: a multicentre retrospective study. Lancet Oncol 26, 136–146 (2025). 10.1016/S1470-2045(24)00599-0

33. Kaczmarzyk, J. R., Saltz, J. H. & Koo, P. K. Explainable AI for computational pathology identifies model limitations and tissue biomarkers. ArXiv (2024).

34 Sarker, I. H. Deep Learning: A Comprehensive Overview on Techniques, Taxonomy, Applications and Research Directions. SN Comput Sci 2, 420 (2021). 10.1007/s42979-021-00815-1

35 Zhang, Y. et al. Attention is all you need: utilizing attention in AI-enabled drug discovery. Brief Bioinform 25 (2023). 10.1093/bib/bbad467

36 Saldanha, O. L. et al. Self-supervised attention-based deep learning for pan-cancer mutation prediction from histopathology. NPJ Precis Oncol 7, 35 (2023). 10.1038/s41698-023-00365-0

37 Jiang, Y. et al. Biology-guided deep learning predicts prognosis and cancer immunotherapy response. Nat Commun 14, 5135 (2023). 10.1038/s41467-023-40890-x

38 Diao, J. A. et al. Human-interpretable image features derived from densely mapped cancer pathology slides predict diverse molecular phenotypes. Nat Commun 12, 1613 (2021). 10.1038/s41467-021-21896-9

39 Abel, J. et al. AI powered quantification of nuclear morphology in cancers enables prediction of genome instability and prognosis. NPJ Precis Oncol 8, 134 (2024). 10.1038/s41698-024-00623-9

40 Amgad, M. et al. A population-level digital histologic biomarker for enhanced prognosis of invasive breast cancer. Nat Med 30, 85–97 (2024). 10.1038/s41591-023-02643-7

41 Rakha, E. A. et al. Prognostic significance of Nottingham histologic grade in invasive breast carcinoma. J Clin Oncol 26, 3153–3158 (2008). 10.1200/JCO.2007.15.5986

42 Elston, C. W. & Ellis, I. O. Pathological prognostic factors in breast cancer. I. The value of histological grade in breast cancer: experience from a large study with long-term follow-up. Histopathology 19, 403–410 (1991). 10.1111/j.1365-2559.1991.tb00229.x

43 Edwards, N. J. et al. The CPTAC Data Portal: A Resource for Cancer Proteomics Research. J Proteome Res 14, 2707–2713 (2015). 10.1021/pr501254j

44 Peikari, M., Salama, S., Nofech-Mozes, S. & Martel, A. L. Automatic cellularity assessment from post-treated breast surgical specimens. Cytometry A 91, 1078–1087 (2017). 10.1002/cyto.a.23244

45 Parker, J. S. et al. Supervised risk predictor of breast cancer based on intrinsic subtypes. J Clin Oncol 27, 1160–1167 (2009). 10.1200/JCO.2008.18.1370

46 Graham, S. et al. Hover-Net: Simultaneous segmentation and classification of nuclei in multi-tissue histology images. Med Image Anal 58, 101563 (2019). 10.1016/j.media.2019.101563

47 Liu, S. et al. A panoptic segmentation dataset and deep-learning approach for explainable scoring of tumor-infiltrating lymphocytes. NPJ Breast Cancer 10, 52 (2024). 10.1038/s41523-024-00663-1

48 Prat, A. et al. Prognostic significance of progesterone receptor-positive tumor cells within immunohistochemically defined luminal A breast cancer. J Clin Oncol 31, 203–209 (2013). 10.1200/JCO.2012.43.4134

49 Rakha, E. A. et al. Breast cancer prognostic classification in the molecular era: the role of histological grade. Breast Cancer Res 12, 207 (2010). 10.1186/bcr2607

50 Desmedt, C. et al. Biological processes associated with breast cancer clinical outcome depend on the molecular subtypes. Clin Cancer Res 14, 5158–5165 (2008). 10.1158/1078-0432.CCR-07-4756

51. Neidlinger, P., et al. A deep learning framework for èicient pathology image analysis. arXiv (2025).

52 Schuiveling, M. et al. A novel dataset for nuclei and tissue segmentation in melanoma with baseline nuclei segmentation and tissue segmentation benchmarks. Gigascience 14 (2025). 10.1093/gigascience/giaf011

53 Ensenyat-Mendez, M. et al. Current Triple-Negative Breast Cancer Subtypes: Dissecting the Most Aggressive Form of Breast Cancer. Front Oncol 11, 681476 (2021). 10.3389/fonc.2021.681476

54 Xiong, N., Wu, H. & Yu, Z. Advancements and challenges in triple-negative breast cancer: a comprehensive review of therapeutic and diagnostic strategies. Front Oncol 14, 1405491 (2024). 10.3389/fonc.2024.1405491

55 He, K. M., Zhang, X. Y., Ren, S. Q. & Sun, J. Deep Residual Learning for Image Recognition. Proc Cvpr Ieee, 770–778 (2016). 10.1109/Cvpr.2016.90

56 Gillies, S. & others, a. Shapely: manipulation and analysis of geometric objects. (2007).

