## Supplementary Figures for "Pathologist-interpretable breast cancer subtyping and stratification from AI-inferred nuclear features"

### **SUPPORTING FIGURES**

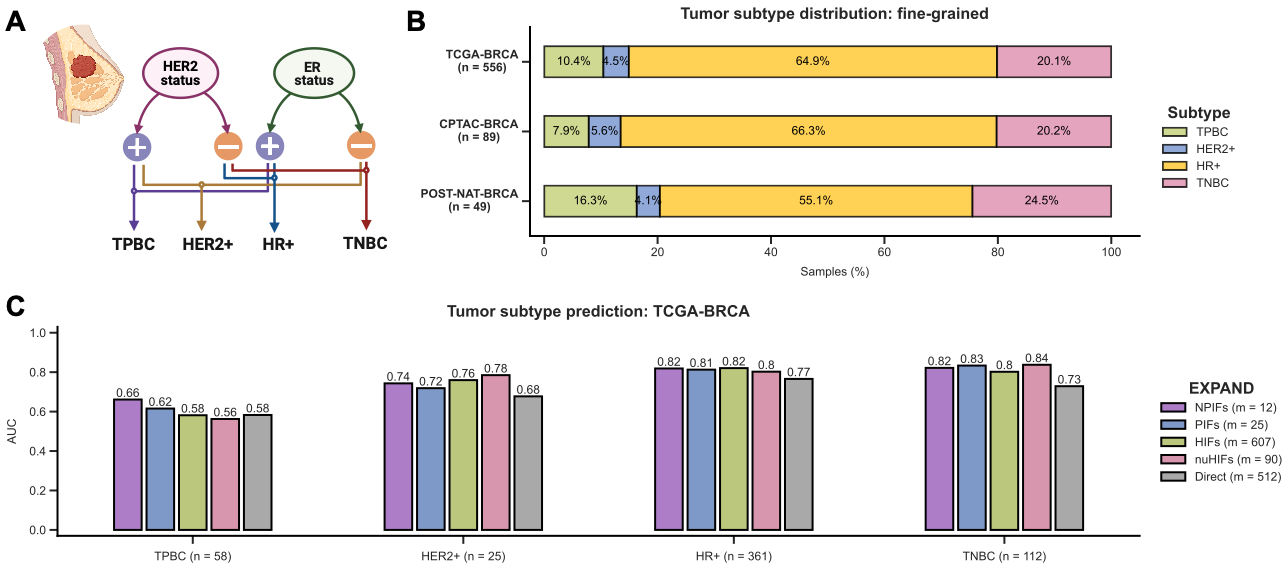

**Figure S1: Pathologist-interpretable features (PIFs) and nuclear PIFs (NPIFs) identify the fine-grained tumor subtypes with high accuracy while maintaining interpretability.**

**A.** Breast tumor fine-grained four tumor subtype definition based on the statuses of HER2 (human epidermal growth factor receptor 2) and ER (estrogen receptor).

**B***.* Distribution of fine-grained tumor subtypes across the histopathology cohorts used for analysis.

**C.** Performance comparison for predicting the fine-grained tumor subtypes in TCGA-BRCA (*n* = 556) using five feature sets: NPIFs, PIFs, HIFs, nuHIFs and direct features. ‘*m*’ denotes the feature set size. AUC stands for the area under the receiver operating characteristics curve.

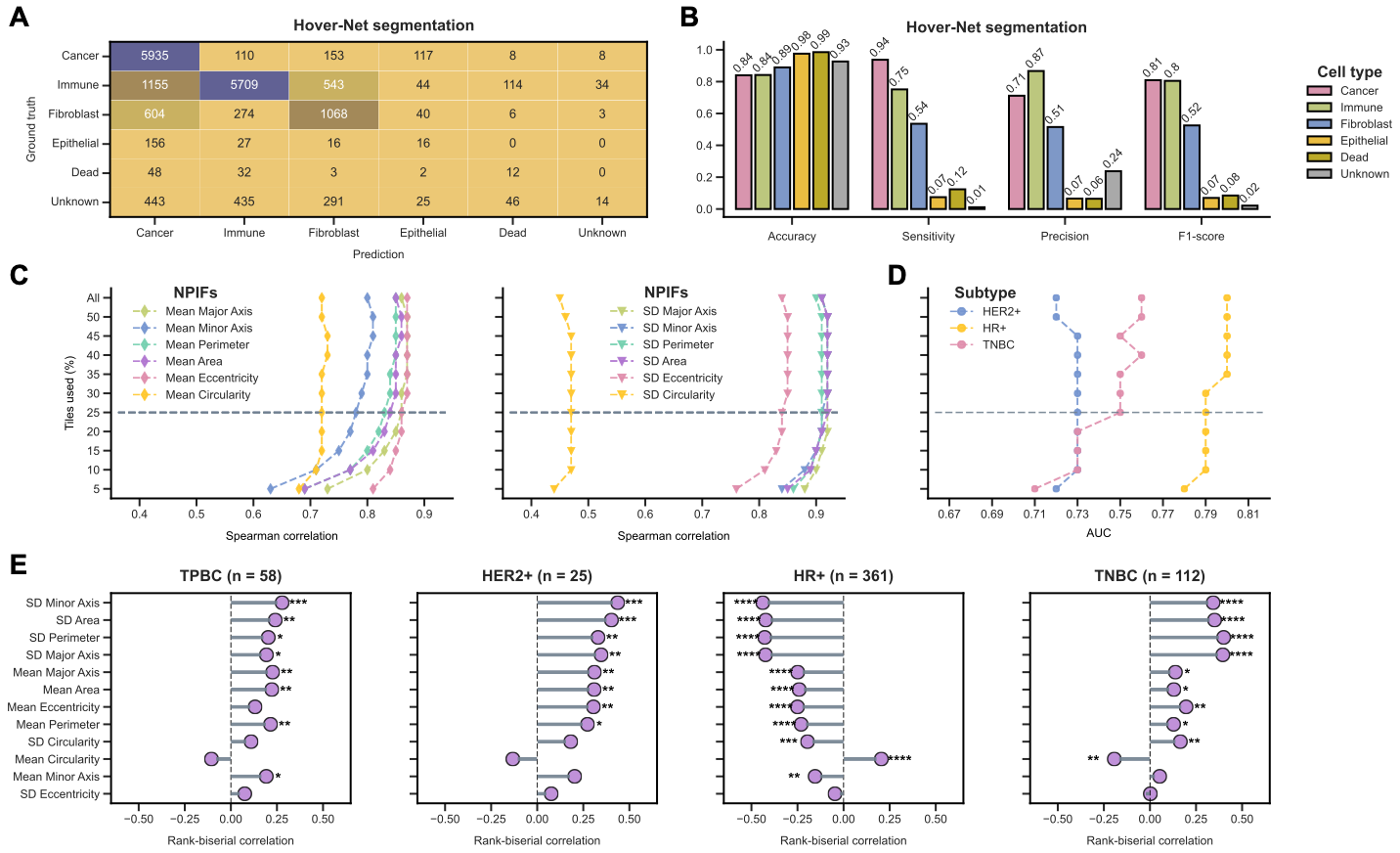

**Figure S2: EXPAND facilitates the robust extraction of NPIFs from whole-slide images.**

**A-B.** Hover-Net nuclei classification performance on the PanopTILs dataset (n = 151), depicted by confusion matrix (**A**) and per-class performance scores (**B**). Prediction and ground truth denote the Hover-Net identified nucleus types and the pathologist-annotated nucleus types, respectively.

**C.** Concordance between the EXPAND-selected NPIFs (from HIFs and nuHIFs) with EXPAND-extracted NPIFs computed from all tiles vs. the top 50% to 5% of the tumor-enriched tiles (gradual decrease of 5%) in TCGA-BRCA (*n* = 556). The gray dotted line indicates the optimal correlation.

**D.** Performance comparison for tumor subtype prediction using the EXPAND-extracted NPIFs computed from all tiles vs. the top 50% to 5% of the tumor-enriched tiles (gradual decrease of 5%) in TCGA-BRCA (*n* = 556). The gray dotted line denotes the optimal performance. AUC stands for the area under the receiver operating characteristics curve.

**E.** Associations of the EXPAND-extracted NPIFs with the fine-grained tumor subtypes in TCGA-BRCA, computed by using two-sided Wilcoxon rank-sum tests (****, ***, ** and * denote P ≤ 0.0001, P ≤ 0.001, P ≤ 0.01 and P ≤ 0.05), where the effect size is measured by rank-biserial correlation coefficient. NPIFs are ordered by their mean absolute effect sizes across subtypes in a descending order.

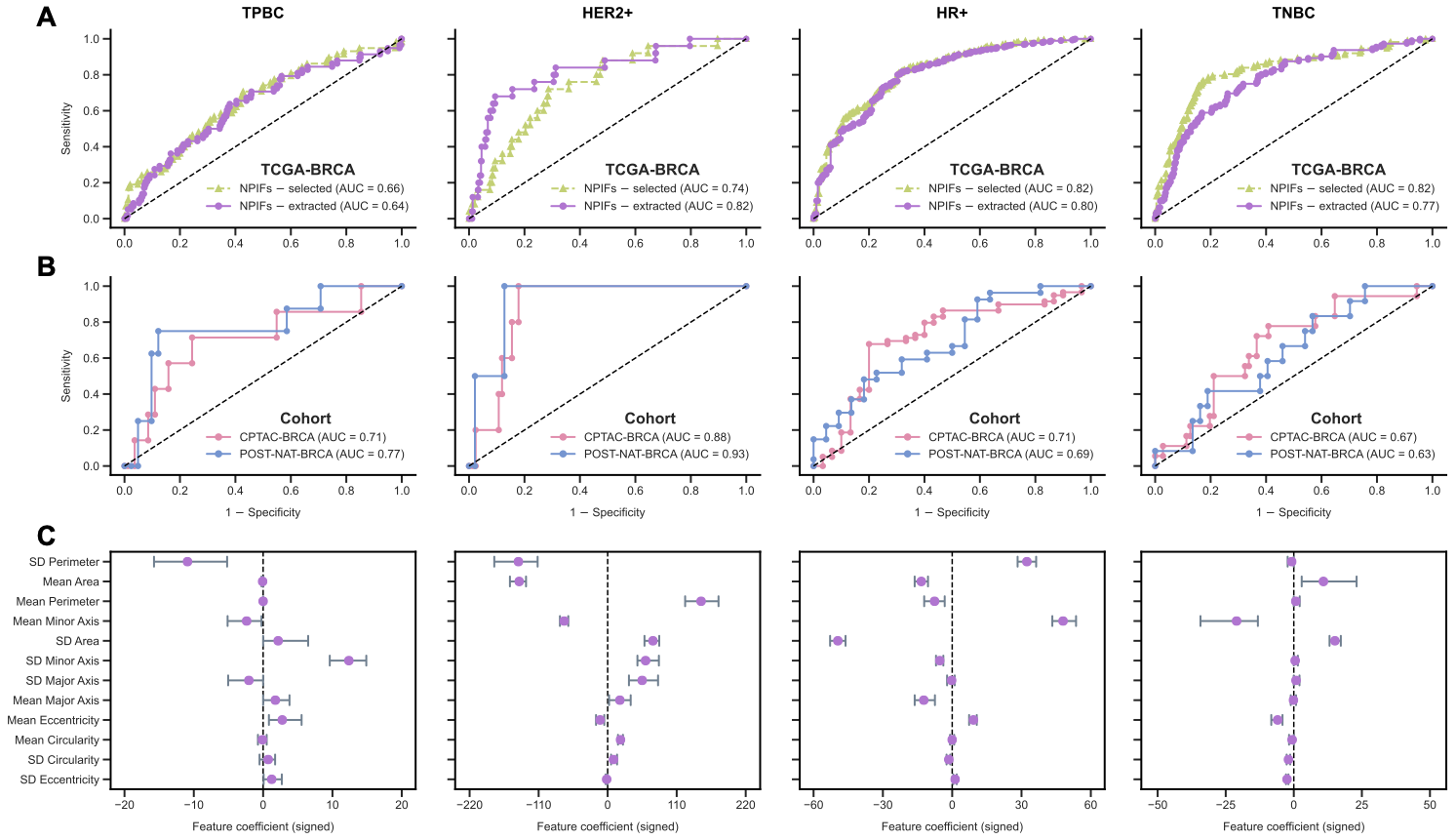

**Figure S3: EXPAND reliably identifies the fine-grained tumor subtypes using both selected and extracted NPIFs, and further generalizes to external cohorts.**

**A.** Fine-grained tumor subtype prediction in TCGA-BRCA (*n* = 556) using EXPAND with the selected and extracted NPIFs. AUC stands for the area under the receiver operating characteristics curve.

**B.** External validation of EXPAND subtype classifiers on the independent cohorts: CPTAC-BRCA (*n* = 89) and POST-NAT-BRCA (*n* = 49).

**C****.** Feature importances (mean with 95% confidence interval; CI) for the NPIFs in fine-grained tumor subtype classification, where higher values indicate high impacts on prediction and *vice versa*, with the sign denoting the direction of impact. NPIFs are ordered by their mean absolute feature importance across subtypes in a descending order.

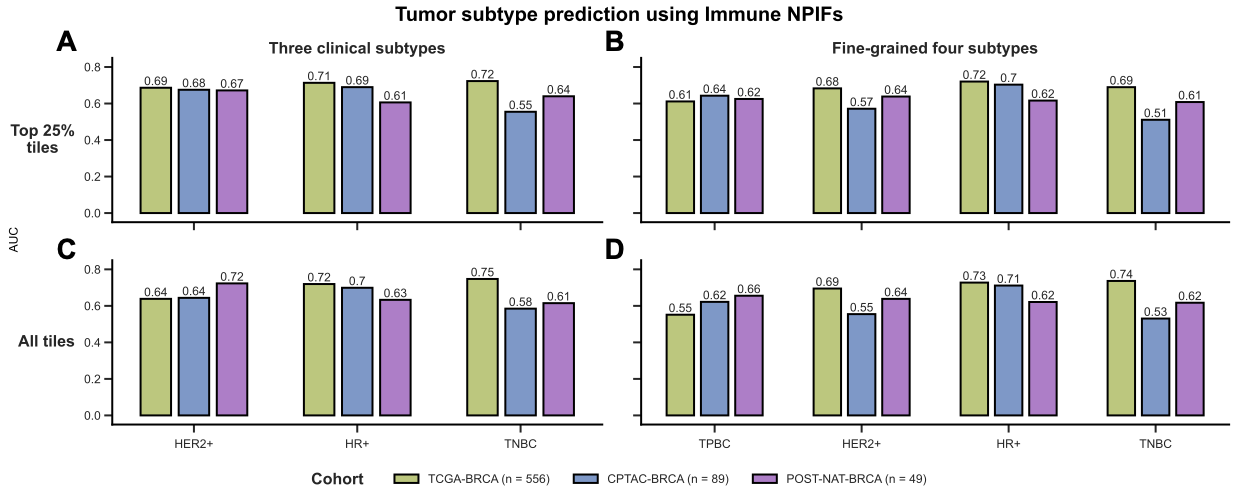

**Figure S4: EXPAND identifies tumor subtypes using the Immune NPIFs.**

**A–B.** Tumor subtype prediction using the Immune NPIFs computed using the top 25% immune-enriched tiles for both the three clinical subtypes (**A**) and the fine-grained four subtypes (**B**). AUC stands for the area under the receiver operating characteristics curve.

**C–D.** Tumor subtype prediction using Immune NPIFs computed using all available tiles with immune nuclei for both the three clinical subtypes (**A**) and the fine-grained four subtypes (**B**).

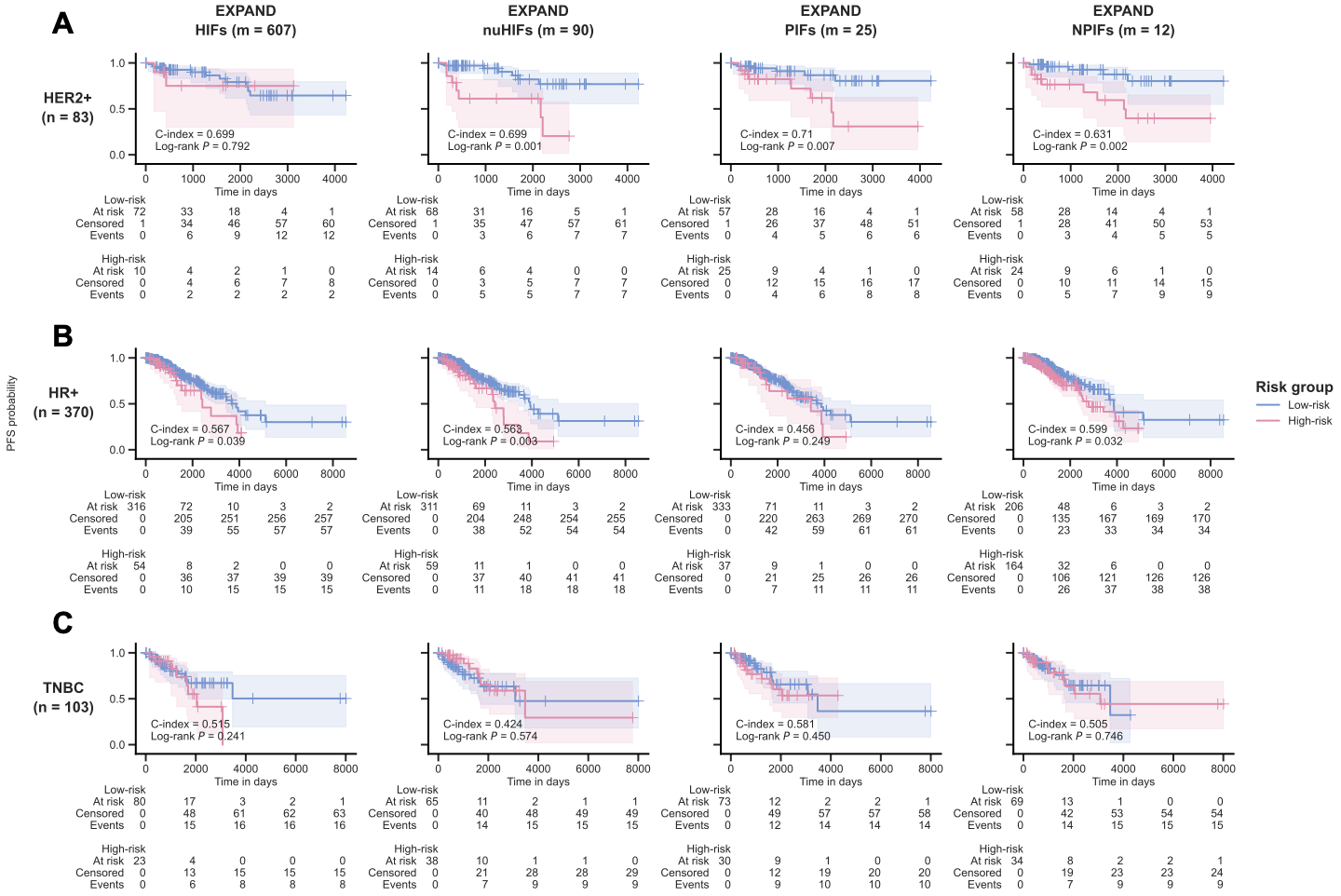

**Figure S5: EXPAND stratifies TCGA-BRCA patient survival within each tumor subtype.**

**A-C.** Kaplan-Meier curves depicting the progression-free survival (PFS) probabilities for TCGA-BRCA patients with HER2+ (**A**), HR+ (**B**) and TNBC (**C**) tumors, stratified using models built with HIFs, nuHIFs, PIFs and NPIFs. Patients were stratified into ‘Low-risk’ and ‘High-risk’ groups by using a fixed threshold of 0.5 on quantile-normalized risk score (using the [10%, 90%] interval). The differences between the two curves were computed by using a Log-rank test (*P* ≤ 0.1).
