## Supplementary Tables for "Pathologist-interpretable breast cancer subtyping and stratification from AI-inferred nuclear features": SupplementaryTableLegends.docx

**Supplementary Table Legends**

**Supplementary Table 1**: Human-interpretable image features (HIFs) with biomarker status for TCGA-BRCA samples.

**Supplementary Table 2**: Nuclear HIFs (nuHIFs) with biomarker status for TCGA-BRCA samples.

**Supplementary Table 3**: Pathologist-interpretable features (PIFs) with biomarker status for TCGA-BRCA samples.

**Supplementary Table 4**: PathAI-derived nuclear pathologist-interpretable features (NPIFs) with biomarker status for TCGA-BRCA samples.

**Supplementary Table 5**: Description of 607 PathAI HIFs.

**Supplementary Tables 6–7**: Description of 90 PathAI nuHIFs.

**Supplementary Table 8**: Description of 25 PIFs.

**Supplementary Table 9**: Description of 12 NPIFs.
